# Predicting Drug Interactions to Unassociated Biomedical Implants Using Machine Learning Techniques and Model Polymers

**DOI:** 10.1101/2020.11.10.374900

**Authors:** Jacob Kerner, Horst von Recum

**Author notes:** Correspondence; Case Western Reserve University.

## Abstract

Affinity based drug delivery mechanisms increase efficacy and minimalize off target effects when compared to non-specific methods due to the localization of drugs within target areas. While this is beneficial for targeted delivery, introduction of foreign polymeric medical devices into the body provide a potential area of localization due to high affinity between administered drugs and polymers. Previous attempts at creating models to predict affinity between small molecule drugs and polymers require a specific model be trained for each individual polymer failing to incorporate input features of both the polymer (host) and small molecule drug (guest). Within, we propose a universal model built using a neural network and quantitative structure activity relationships to predict the binding energy between guest and host molecules using input features. The trained model returned a correlation value, R^2^, of 0.9806 and 0.9958 between predicted and experimental binding affinity for the training and validation sets, respectively. This correlates to a mean absolute error of 0.951 kJ/mol and 0.771 kJ/mol for the training and validation sets, respectively. While limited to the current polymers used to train the model, the dataset can be expanded, and models retrained for further applications.

## 1. Introduction

Polymers have become commonplace in medicine being used for a variety of applications due to their wide and customizable range of mechanical, electrical, chemical, and thermal properties.^1^ Polymers used for the bulk manufacturing of devices comprises only a fraction of current utilization. Applications of polymer biomaterials include but are not limited to; surface coatings, adhesives, and nanocarriers.^2,3^ Due to their prevalence, procedures involving the implantation of devices including some form of polymer material (i.e. drug eluting stents and knee/hip replacements) occur millions of time every year in the United States alone.^4^ Removal of polymer materials currently rely primarily on surgical removal or degradation pathways within the body. It is generally assumed that due to the biocompatibility of polymers, there will be no adverse reactions if polymer materials are not removed.^5^ Therefore, little published information addressing the interactions of polymers with various medications and biomolecules is available.

Poly(lactic-co-glycolic acid), PLGA, is a commonly used biomaterial polymer, showing excellent biocompatibility in devices ranging from sutures to screws to stents.^6^ Recent data has shown that the chemotherapeutic drug doxorubicin, which would typically be unrelated to any PLGA usage, was capable of selective localization in PLGA implants in vivo. Intentional localization of medication, while championed in the drug delivery field (e.g. drug targeting, reloading), can have cytotoxic effects if unwanted depending on the concentrations achieved and medication used. Molecular interactions, such as binding affinity between antibodies and antigens to deliver such cytotoxic drugs has commonly been examined in the pharmaceutical industry for targeted localization of chemotherapy agents.^7–9^ Binding affinity is commonly measured by the change in Gibbs free energy (ΔG) when binding occurs. While interactions between polymers and drug molecules have lower binding affinity when compared to specific antibody/antigen binding (ΔG≈-15kJ/mol compared to ΔG <-50kJ/mol), the potential for unintentional accumulation due to binding affinity interactions cannot be disregarded.^10,11^ Accumulation of medications in biomaterial implants has the potential to not only generate cytotoxic effects in surrounding tissue, but also degrade polymers through side reactions. Degradation of polymers can be attributed to various mechanisms, however all lead to undesirable structure and mechanical properties leading to higher rates of implant failure.^12,13^

Pharmaceutical drug development and discovery commonly uses quantitative structure activity relationships (QSAR) to predict binding between drug molecules and intended targets. QSAR techniques use numerical values to quantify properties of molecules (i.e. hydrophilicity and electronegativity) before inputting into a model or scanning for similar drugs. There is currently well established methods for generating QSAR models using machine learning techniques for drug identification and discovery.^14,15^ Recently applications of QSAR models to materials, aptly named Quantitative Structure Property Relationships (QSPR), have been successful in predicting material properties *in silico*.^16–18^ There have been previous attempts at computational models to predict interactions between polymers and small molecules drugs however, these previous iterations have been limited to one polymer type per model with varying accuracy.^11,19^ A singular model capable of predicting interactions between various polymer subunits and small molecule drugs has currently eluded researchers.

In this work, we propose a novel QSAR strategy using polymers made from cyclodextrins as model polymers. To obtain QSAR descriptors for the selected polymers and small molecule drugs, the open source software PaDEL descriptor was used.^20^ Encoders were used to reduce the dimensionality of the features of the model. Features were passed to an artificial neural network used as a regressor, trained to predict the Gibbs free energy change of binding between the two molecules. Network hyperparameters were optimized by minimizing the error in predicted results. The results of this study is a development of a new QSAR model capable of predicting binding affinity between small molecule drugs and model polymers in a biologically relevant environment.

## 2. Methodology

### 2.1 Data Set

Cyclodextrins (CD) are small cyclical oligosaccharides that have long been explored in the pharmaceutical industry to improve the solubility of hydrophobic drugs.^21,22^ CD molecules have been investigated for use in slow sustained drug delivery applications, exploiting the CD and small molecule drug complexes to slow delivery rates beyond that capable of diffusion alone. While more advantageous to use FDA approved polymers (PLGA), CDs offer a unique advantage in the form of high availability of previously published binding affinity values for various small molecule drugs. There are currently well established data sets for complexations between three CD (alpha, beta, and gamma) and small molecule drugs available for model training. In this work, the data set used for the training and testing of the proposed model were collected from[9]. The cyclodextrin data set consisted of 2000+ binding energies between small molecule drugs and biologics with the three different forms of cyclodextrin polymers. The dataset was split into training and test sets with a 95%/5% respectively.

Prior to descriptor calculation, combinations in buffer solutions outside of biological relevant ranges were removed from the data set. Binding affinity energies that were not measured in a water solution were immediately disqualified. The pH of the solution was restricted to a range of 6.9 to 7.7, similar to that found within the body and blood (pH of 7.4), to ensure charges located on functional groups remained constant. Temperatures during measurement were restricted to 273K to 315K due to most measurements being made at STP or room temperature. After removal of nonbiologically relevant solutions, 725 model polymer/small molecule drug combinations remained.

### 2.2 Molecular Descriptors

PaDEL descriptor is capable of calculating 1444 2D and 431 3D descriptors for each molecule. All molecular files used within PaDEL were collected from PubChem utilizing the python library and a custom script.^23^ These descriptors can be classified into several categories including; constitutional descriptors, molecular properties, structural descriptors, geometric descriptors, Eigenvalues, and molecular fingerprints. After calculation, each model polymer/molecule combination contained approximately 3650 molecular descriptors.

### 2.3 Artificial Neural Networks

QSAR models using artificial neural networks (ANN) have been proven successful in various previous studies.^24,25^ ANN’s consist of three major layers; an input, hidden layers consisting of interconnected nodes called neurons, and an output layer. These neurons act in similar ways to their biological counterparts as they are summation nodes. Inputs are fed into the node, summed, multiplied by a scalar weight value, and then passed to the next neuron (Figure 1A). Activation functions (*f)* as well as biases (*b)* can be added for further modifications. Models are trained by inputting data sets into the model and calculating the outputs. The output is compared to the target before adjusting model weights and biases to determine the optimal values (Figure 1B). When the entire dataset has been cycled through, this is known as an epoch.

**Figure 1.**
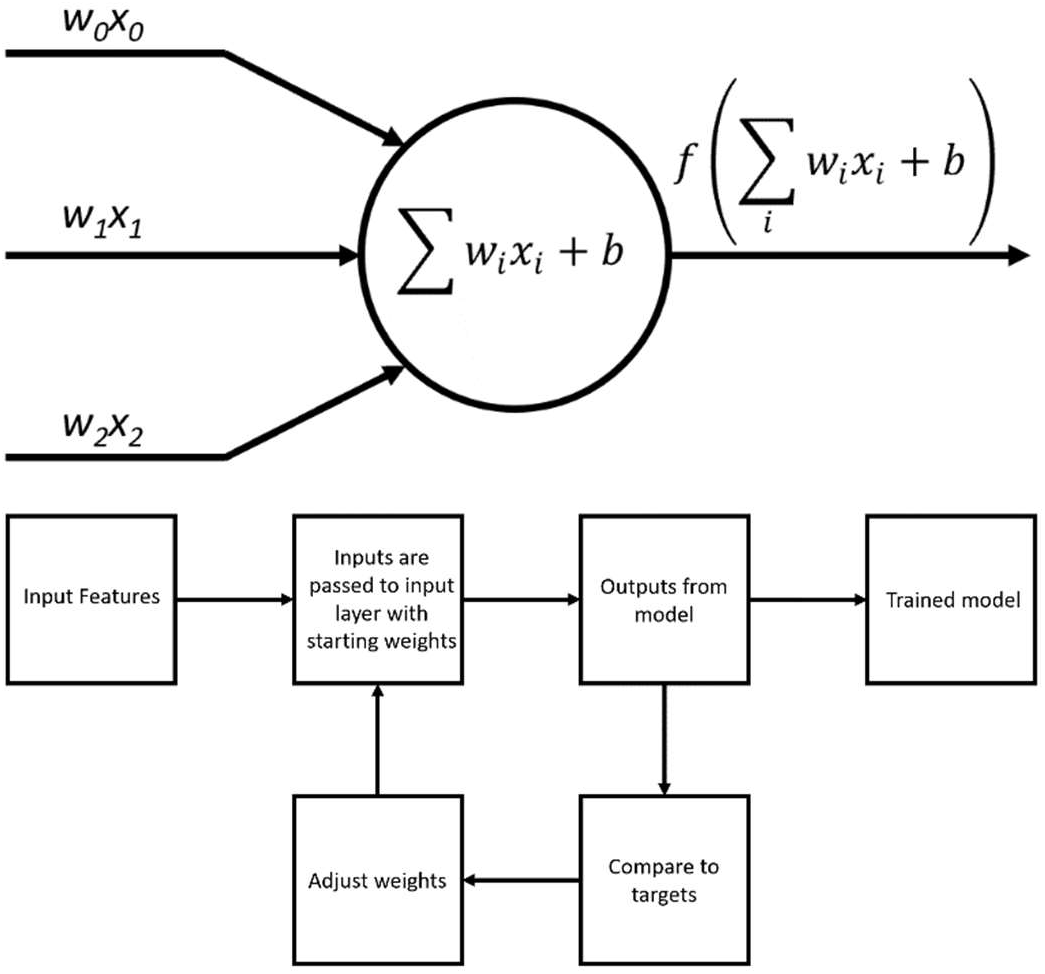
A (Top). Overview of a neuron used as the basic computational unit in an ANN. B (Bottom). Repetitive training method applied to ANN to reduce error.

### 2.4 Feature Manipulation and Dimensionality Reduction

Due to the large number of descriptors, those with low variability were removed reducing the total number of input features to 1433. Prior to feature extraction, each descriptor was normalized to prevent features with smaller magnitudes becoming neglected. To further reduce the dimensionality of the model input features, autoencoders were implemented.^26^ Autoencoders are a form of unsupervised learning structure consisting of two parts: an encoder and a decoder (Figure 2). Encoders are used for mapping features to a lower dimensional representation while decoders reconstruct the original features from the lower dimensional representation. Encoded data can be represented by:

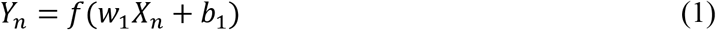

where *Y*_*n*_ is the encoded data vector, *X*_*n*_ is the input feature vector, *w*_*1*_ is the weight matrix of the encoder, *b*_*1*_ is the bias vector of the encoder, and *f* is the encoding function. The decoder output can be represented by:

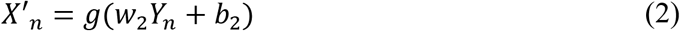

where *X*’_*n*_ is the reconstructed data vector, *w*_*2*_ is the weight matrix of the decoder, *b*_*2*_ is the bias vector of the decoder, and *g* is the decoding function.

**Figure 2.**
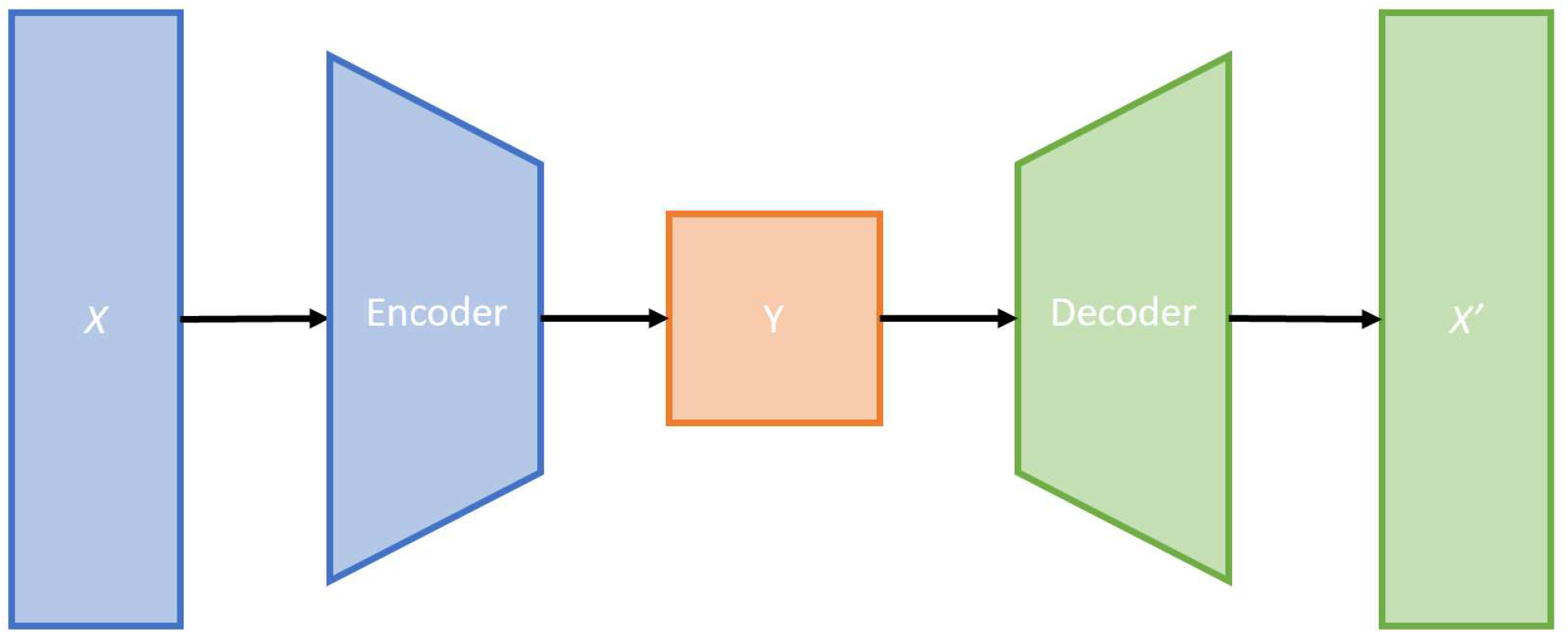
Structure of autoencoder used to reduce dimensionality of input features.

It was determined 250 input features each for the model polymer and the small molecule drug would be sufficient, a dimensionality reduction of 65.2%. The hidden layers of the encoder and decoder were optimized by randomly varying the size of the hidden layer to minimize the reconstructed feature error:

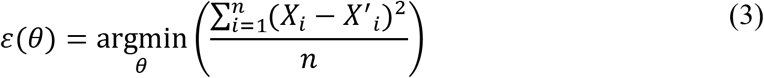

where *ε* is the error, *θ* is the hyperparameter setting (hidden layer size), and *n* is the size of the feature vector. Weights for connections, *w*_*1*_ and *w*_*2*_, are trained using data and minimizing the given error function. Each autoencoder consisted of a single fully connected hidden layer consisting of *θ* neurons with linear activation trained for 1000 epochs using the ADAM optimizer. After training, only the encoder will be used in the final combined model.

### 2.5 Proposed Model Design

A model to predict the Gibbs free energy change of binding between the model polymers and small molecule drugs was created consisting of two input layers and a singular output. The model consisted of three distinct artificial neural networks, two encoders and a dense neural network which will be trained for binding affinity prediction (Figure 3). The two inputs layers, one for each set of features (model polymer and drug molecule) were fed to the trained feature encoders. Outputs from the encoders were combined prior to being passed into the single input layer for the dense neural network used for affinity prediction.

**Figure 3.**
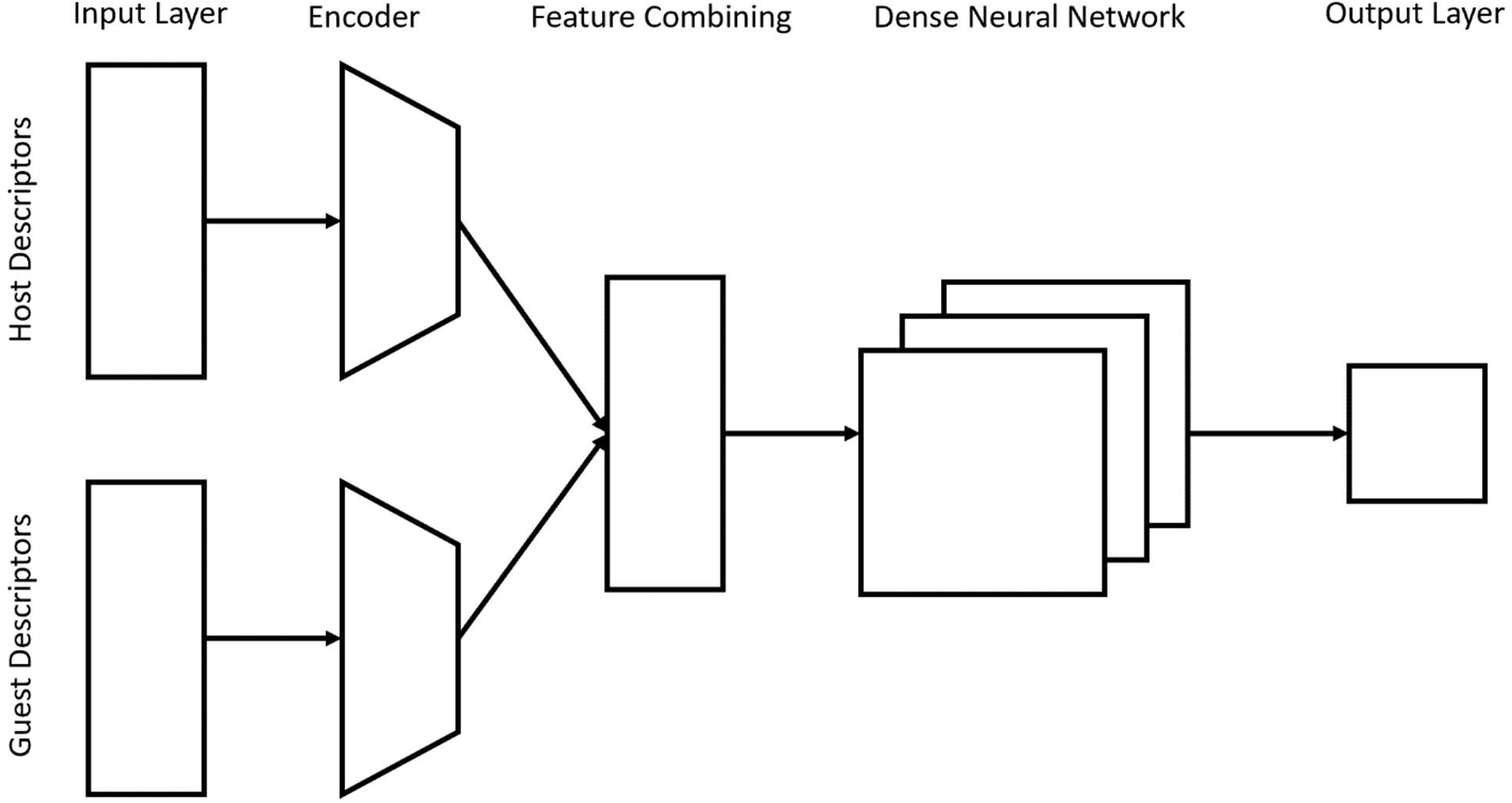
Schematic for proposed model design.

Hyperparameters are model settings (i.e. number of hidden layers and number of neurons per layer) the model cannot learn through training. Three hyperparameters were chosen to vary: the number of layers (1-3), the number of neurons per layer (1-300), and the activation of the layer (none, relu, linear, tanh, sigmoid). To determine the optimal hyperparameter for the dense neural network, network design was optimized using a custom script which determined the loss of each trained network by testing various hyperparameter settings. The loss function minimized while training each model can be described as:

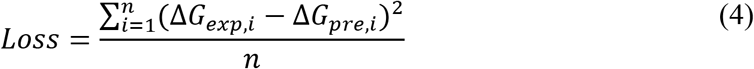

where Δ*G*_*exp*_ is the experimentally reported binding affinity and Δ*G*_*pre*_ is the predicted binding affinity. Hyperparameters combinations were randomized into a matrix before being sliced into vectors. Each vector of hyperparameter settings were used to train each model for 100 epochs using the ADAM optimizer. Hyperparameter optimization function can be described by combing equation 3 and equation 4:

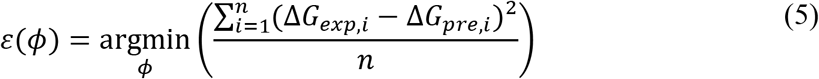

where *ϕ* is the vector of hyperparameter settings. Hyperparameter settings that returned the lowest loss value were for the model were saved and the trained model itself was saved for later verification on the test data set.

### 3. Results and Discussion

After 1000 epochs, the polymer and small molecule drug autoencoder returned loss values of 0.0904 and 0.0541 correlating to a mean absolute error of reconstructed features from encoded data of 0.0994 and 0.0887, respectively. To further verify the accurate mapping of features, a correlation coefficient, R^2^ was used to compare reconstructed features to original features. Larger error in the polymer dataset is more than likely due to the significantly lower number of samples in the training set. Due to the low error value in reconstructed features and high correlation coefficients, encoders were deemed sufficient and were implemented as input layers in the model.

The optimal design of the model determined by our script consisted of a total of three hidden layers; two with 100 neurons, hyperbolic tangent activation and one with one neuron, linear activation (Figure 4). This model returned a mean absolute error in predictions of 0.951 on the training set. To further verify the model, binding affinities of a polymer/drug test set taken from the original dataset as a validation set were predicted. The validation set returned a mean absolute error of 0.771. Correlation coefficients were calculated along with training metrics for each dataset, training set and test set which can be found in Table 1. Experimentally determined binding affinity and predicted binding affinity was plotted and high correlation can be seen in both the training and test datasets (Figure 5). Values of experimental and predicted binding affinity (ΔG) for each cyclodextrin polymer can be seen in tables 2–4.

**Table 1.** Training metrics used to evaluate the accuracy of various data sets.

| Data Set | Mean Absolute Error (kJ/mol) | Correlation Coefficient ($R^2$) |
| --- | --- | --- |
| **Training** | 0.951 | 0.9806 |
| **Validation** | 0.771 | 0.9958 |

**Table 2.** Experimental and predicted free energy change for alpha cyclodextrin.

| MOLECULE NAME | EXPERIMENTAL<br>(KJ/MOL) | PREDICTED<br>(KJ/MOL) | ABSOLUTE ERROR<br>(KJ/MOL) |
| --- | --- | --- | --- |
| 3-CHLOROPHENYL ACETATE | -12.9 | -12.67 | 0.23 |
| 3-ETHYLPHENOL | -10.84 | -10.98 | 0.14 |
| CYCLOOCTANOL | -12.84 | -13.57 | 0.73 |
| P-TOLYL ACETATE | -11.19 | -11.26 | 0.07 |
| (1R,2R)-(-)-PSEUDOEPHEDRINE | -7.25 | -7.92 | 0.67 |
| (1R,2S)-(-)-EPHEDRINE | -7.02 | -7.86 | 0.84 |
| (1S,2R)-(+)-EPHEDRINE | -7.17 | -7.86 | 0.69 |
| (1S,2S)-(+)-PSEUDOEPHEDRINE | -7.41 | -7.92 | 0.51 |
| (2,5-DIMETHOXYPHENETHYL)AMMONIUM | -8.81 | -8.80 | 0.01 |
| (3,4-DIMETHOXYPHENETHYL)AMMONIUM | -5.1 | -6.61 | 1.51 |
| (3-METHOXYPHENETHYL)AMMONIUM | -7.73 | -7.97 | 0.24 |
| (3-PHENYLPROPYL)AMMONIUM | -7.85 | -9.23 | 1.38 |
| (4-METHOXYPHENETHYL)AMMONIUM | -9.39 | -9.10 | 0.29 |
| (4-METHYLPHENETHYL)AMMONIUM | -10.17 | -9.53 | 0.64 |
| (R)-(-)-2-BUTANOL | -7.9 | -7.53 | 0.37 |
| (R)-(-)-2-HEPTANOL | -16.4 | -16.62 | 0.22 |
| (R)-(-)-2-HEXANOL | -14.5 | -14.90 | 0.40 |
| (R)-(-)-2-PENTANOL | -12 | -12.03 | 0.03 |
| (R)-(-)-PHENYLEPHRINE | -3.5 | -4.03 | 0.53 |
| (S)-(+)-2-BUTANOL | -7.9 | -7.53 | 0.37 |
| (S)-(+)-2-HEPTANOL | -16.5 | -16.62 | 0.12 |
| (S)-(+)-2-HEXANOL | -14.5 | -14.90 | 0.40 |
| (S)-(+)-2-PENTANOL | -12 | -12.03 | 0.03 |
| 1,10-DECANEDIOL | -22 | -22.12 | 0.12 |
| 1,2-BUTANEDIOL | -6.3 | -7.02 | 0.72 |
| 1,2-ETHANEDIOL | 0.57 | -0.05 | 0.62 |
| 1,2-HEXANEDIOL | -12.9 | -12.57 | 0.33 |
| 1,2-PENTANEDIOL | -10.8 | -11.12 | 0.32 |
| 1,2-PROPANEDIOL | -2.7 | -2.95 | 0.25 |
| 1,3-BUTANEDIOL | -5.7 | -7.32 | 1.62 |
| 1,3-PROPANEDIOL | -3.03 | -3.35 | 0.32 |
| 1,4-BUTANEDIOL | -5.2 | -6.70 | 1.50 |
| 1,4-DIBROMOBENZENE | -17.07 | -17.49 | 0.42 |
| 1,5-HEXANEDIOL | -8.8 | -11.27 | 2.47 |
| 1,5-PENTANEDIOL | -8.56 | -9.60 | 1.04 |
| 1,6-HEXANEDIOL | -11.25 | -12.38 | 1.13 |
| 1,7-HEPTANEDIOL | -14.3 | -15.59 | 1.29 |
| 1,8-OCTANEDIOL | -17.6 | -17.64 | 0.04 |
| 1,9-NONANEDIOL | -20.3 | -20.51 | 0.21 |
| 11-AMINOUNDECANOIC ACID | -19 | -18.20 | 0.80 |
| 1-ADAMANTANEACETATE | -12.97 | -13.66 | 0.69 |
| 1-ADAMANTANECARBOXYLATE | -12.3 | -12.53 | 0.23 |
| 1-ADAMANTYLAMMONIUM | -13.9 | -14.37 | 0.47 |
| 1-BENZYLIMIDAZOLE | -10.15 | -12.10 | 1.95 |
| 1-BUTANOL | -10.95 | -10.87 | 0.08 |
| 1-BUTYLIMIDAZOLE | -12.77 | -12.70 | 0.07 |
| 1-CHLORO-4-NITROBENZENE | -10.45 | -11.45 | 1.00 |
| 1-HEPTANOL | -19.18 | -19.77 | 0.59 |
| 1-HEXANESULFONATE | -11.73 | -12.38 | 0.65 |
| 1-HEXANOL | -16.84 | -16.76 | 0.08 |
| 1-HYDROXY-2,2,5,5-TETRAMETHYLPYRROLIDINE-3-CARBOXAMIDE | -3 | -3.03 | 0.03 |
| 1-NAPHTHALENEACETATE | -16.8 | -17.37 | 0.57 |
| 1-OCTANOL | -21.69 | -21.71 | 0.02 |
| 1-PENTANOL | -14.33 | -15.59 | 1.26 |
| 1-PROPANOL | -7.82 | -8.50 | 0.68 |
| 2,2-DIMETHYL-1-PROPANOL | -8.39 | -8.52 | 0.13 |
| 2,5-HEXANEDIOL | -8.1 | -10.08 | 1.98 |
| 2,7-NAPHTHALENEDISULFONATE | -5.6 | -6.36 | 0.76 |
| 2-AMINOBENZOIC ACID | -28.5 | -27.48 | 1.02 |
| 2-BUTANOL | -8.1 | -7.53 | 0.57 |
| 2-HEXANOL | -14.56 | -14.90 | 0.34 |
| 2-METHYL-1-BUTANOL | -11.64 | -11.71 | 0.07 |
| 2-METHYL-1-PROPANOL | -8.22 | -9.26 | 1.04 |
| 2-METHYL-2-BUTANOL | -8.73 | -8.20 | 0.53 |
| 2-METHYL-2-PROPANOL | -3.65 | -3.43 | 0.22 |
| 2-METHYLCYCLOHEXANONE | -8.4 | -10.50 | 2.10 |
| 2-METHYLPROPYLAMINE | -11.5 | -10.22 | 1.28 |
| 2-NAPHTHALENESULFONATE | -14.6 | -15.18 | 0.58 |
| 2-NITROPHENOL | -20.9 | -21.83 | 0.93 |
| 2-NORBORNANEACETATE | -12.55 | -13.26 | 0.71 |
| 2-OCTANOL | -17.98 | -18.23 | 0.25 |
| 2-PENTANOL | -12.16 | -12.03 | 0.13 |
| 2-PROPANOL | -3.94 | -4.07 | 0.13 |
| 3,3-DIMETHYL-2-BUTANOL | -7.42 | -7.66 | 0.24 |
| 3,5-DIBROMOPHENOL | -17.81 | -18.51 | 0.70 |
| 3,5-DICHLOROPHENOL | -14.44 | -15.04 | 0.60 |
| 3,5-DIMETHOXYPHENOL | -6.79 | -7.20 | 0.41 |
| 3-[(4-HYDROXYPHENYL)AZO]BENZOATE | -21.23 | -21.89 | 0.66 |
| 3-AMINOBENZOIC ACID | -9.9 | -12.22 | 2.32 |
| 3-AMINOMETHYL-PROXYL | -3.9 | -4.09 | 0.19 |
| 3-BROMOPHENOL | -14.04 | -16.47 | 2.43 |
| 3-BROMOPHENYL ACETATE | -15.98 | -16.55 | 0.57 |
| 3-CHLOROPHENOL | -12.39 | -13.27 | 0.88 |
| 3-CYANOPHENYL ACETATE | -9.93 | -11.53 | 1.60 |
| 3-ETHOXYPHENOL | -9.65 | -10.49 | 0.84 |
| 3-ETHOXYPHENYL ACETATE | -9.87 | -10.27 | 0.40 |
| 3-ETHYLPHENYL ACETATE | -10.9 | -13.18 | 2.28 |
| 3-FLUOROPHENYL ACETATE | -6.56 | -8.16 | 1.60 |
| 3-HEXANOL | -12.5 | -13.44 | 0.94 |
| 3-HYDROXYBENZOIC ACID | -15.3 | -15.55 | 0.25 |
| 3-IODOPHENOL | -17.81 | -18.44 | 0.63 |
| 3-IODOPHENYL ACETATE | -18.95 | -19.90 | 0.95 |
| 3-METHOXYPHENOL | -5.88 | -8.89 | 3.01 |
| 3-METHYL-1-BUTANOL | -10.67 | -10.58 | 0.09 |
| 3-METHYL-2-BUTANOL | -7.25 | -7.13 | 0.12 |
| 3-METHYLBENZOIC ACID | -13.8 | -14.00 | 0.20 |
| 3-METHYLPHENYL ACETATE | -8.39 | -10.90 | 2.51 |
| 3-N-BUTYLPHENOL | -16.56 | -17.94 | 1.38 |
| 3-N-BUTYLPHENYL ACETATE | -16.27 | -16.48 | 0.21 |
| 3-NITROANILINE | -10.7 | -12.19 | 1.49 |
| 3-NITROPHENOL | -12.1 | -14.07 | 1.97 |
| 3-N-PROPYLPHENYL ACETATE | -13.82 | -14.80 | 0.98 |
| 3-PENTANOL | -11.08 | -11.50 | 0.42 |
| 3-PHENYLPROPIONATE | -8.54 | -10.92 | 2.38 |
| 3-PHENYLPROPIONIC ACID | -17.6 | -16.99 | 0.61 |
| 3-PROPYLPHENOL | -14.33 | -15.64 | 1.31 |
| 3-TERT-BUTYLPHENOL | -8.33 | -8.57 | 0.24 |
| 4-[(4-HYDROXYPHENYL)AZO]BENZOATE | -20.72 | -21.67 | 0.95 |
| 4-AMINOBENZOIC ACID | -16 | -17.25 | 1.25 |
| 4-BENZYLPIPERIDINE | -9.43 | -10.95 | 1.52 |
| 4-BROMOPHENOL | -16.44 | -17.26 | 0.82 |
| 4-CHLOROPHENOL | -13.99 | -14.44 | 0.45 |
| 4-CHLOROPHENYL ACETATE | -10.28 | -11.18 | 0.90 |
| 4-ETHOXYPHENOL | -10.67 | -11.87 | 1.20 |
| 4-ETHYLPHENOL | -12.16 | -14.07 | 1.91 |
| 4-FLUOROPHENOL | -5.2 | -6.80 | 1.60 |
| 4-HYDROXYBENZALDEHYDE | -9.99 | -11.08 | 1.09 |
| 4-HYDROXYBENZOIC ACID | -17.52 | -18.32 | 0.80 |
| 4-HYDROXYCINNAMIC ACID | -18.84 | -20.03 | 1.19 |
| 4-iodoaniline | -19.7 | -19.65 | 0.05 |
| 4-iodoanilinium | -17.3 | -17.67 | 0.37 |
| 4-iodophenol | -19.01 | -19.54 | 0.53 |
| 4-METHOXYPHENOL | -9.08 | -9.72 | 0.64 |
| 4-METHYL-2-PENTANOL | -9.82 | -9.40 | 0.42 |
| 4-N-AMYLPHENOL | -19.18 | -20.45 | 1.27 |
| 4-N-BUTYLPHENOL | -18.61 | -18.74 | 0.13 |
| 4-NITROANILINE | -15.47 | -15.45 | 0.02 |
| 4-NITROBENZOIC ACID | -12.97 | -13.84 | 0.87 |
| 4-NITROPHENOL | -12.55 | -14.09 | 1.54 |
| 4-NITROPHENOLATE | -18.7 | -19.47 | 0.77 |
| 4-NITROPHENYL ACETATE | -10.46 | -11.05 | 0.59 |
| 4-NITROPHENYL BETA-D-XYLOSIDE | -11.6 | -12.34 | 0.74 |
| 4-NITROPHENYL-BETA-D-GALACTOSIDE | -10.7 | -11.73 | 1.03 |
| 4-NITROPHENYL-BETA-D-GLUCOSIDE | -10.7 | -11.73 | 1.03 |
| 4-O-METHYLDOPAMINE | -4.1 | -5.28 | 1.18 |
| 4-PROPYLPHENOL | -15.7 | -16.82 | 1.12 |
| 4-TERT-BUTYLPHENOL | -10.96 | -11.58 | 0.62 |
| 5-METHYLRESORCINOL | -6.7 | -7.50 | 0.80 |
| 6-AMINOHEXANOIC ACID | -7.6 | -7.82 | 0.22 |
| 6-HEPTENOATE | -15.05 | -14.42 | 0.63 |
| 8-AMINOOCCTANOIC ACID | -10.7 | -12.89 | 2.19 |
| ACETIC ACID | -21.8 | -21.78 | 0.02 |
| ACETONITRILE | -4.28 | -3.79 | 0.49 |
| ANISOLE | -12.22 | -13.01 | 0.79 |
| ANTHRACENE | -10.7 | -24.65 | 13.95 |
| BENZ[A]ANTHRACENE | -11.1 | -29.50 | 18.40 |
| BENZALDEHYDE | -19.35 | -17.63 | 1.72 |
| BENZENE | -14.35 | -13.58 | 0.77 |
| BENZO[A]PYRENE | -12.8 | -29.75 | 16.95 |
| BENZOATE | -5.8 | -10.46 | 4.66 |
| BENZOIC ACID | -17.15 | -17.88 | 0.73 |
| BENZYL ALCOHOL | -7.59 | -10.78 | 3.19 |
| BROMOBENZENE | -15.47 | -15.75 | 0.28 |
| BUTANOIC ACID | -12.15 | -12.12 | 0.03 |
| BUTYLAMINE | -11.6 | -10.89 | 0.71 |
| BUTYLAMMONIUM | -6.59 | -8.22 | 1.63 |
| CARBON TETRACHLORIDE | -9.25 | -9.06 | 0.19 |
| CHLORCYCLIZINE | -17.6 | -17.67 | 0.07 |
| CHLOROFORM | -9.25 | -9.34 | 0.09 |
| CIS-1,2-CYCLOHEXANEDIOL | -6.9 | -7.35 | 0.45 |
| CYCLOBUTANOL | -9.08 | -8.86 | 0.22 |
| CYCLOHEPTANOL | -10.5 | -11.26 | 0.76 |
| CYCLOHEXANECARBOXYLIC ACID | -9.7 | -10.47 | 0.77 |
| CYCLOHEXANOL | -10.33 | -9.05 | 1.28 |
| CYCLOHEXANONE | -7.7 | -7.84 | 0.14 |
| CYCLOPENTANOL | -9.48 | -10.33 | 0.85 |
| D-FRUCTOSE | -9.8 | -9.92 | 0.12 |
| D-GALACTOSE | -0.68 | -9.38 | 8.70 |
| D-GLUCOSE | -15.1 | -9.38 | 5.72 |
| DIETHYLAMINE | -8.8 | -9.28 | 0.48 |
| DI-TERT-BUTYL NITROXIDE | -4.5 | -4.32 | 0.18 |
| D-MANNOSE | -10.1 | -9.38 | 0.72 |
| D-NORLEUCINE | -9.5 | -9.48 | 0.02 |
| D-NORVALINE | -6.2 | -6.94 | 0.74 |
| D-XYLOSE | -9 | -9.08 | 0.08 |
| ETHANOL | -4.28 | -3.82 | 0.46 |
| ETHYLBENZENE | -11.65 | -13.61 | 1.96 |
| FLUOROBENZENE | -8.73 | -9.09 | 0.36 |
| FLURBIPROFEN | -10.5 | -12.06 | 1.56 |
| FORMIC ACID | -3.5 | -4.18 | 0.68 |
| FUMARIC ACID | -18.5 | -14.19 | 4.31 |
| GLUTARIC ACID | -14.1 | -14.05 | 0.05 |
| HEPTANEDIOIC ACID | -16.4 | -16.76 | 0.36 |
| HEPTANOATE | -16.7 | -16.17 | 0.53 |
| HEXANEDIOIC ACID | -14.3 | -14.53 | 0.23 |
| HEXANOATE | -14.1 | -13.20 | 0.90 |
| HEXANOIC ACID | -16.59 | -15.66 | 0.93 |
| HEXYL BETA-D-GLUCOPYRANOSIDE | -16.69 | -16.90 | 0.21 |
| HEXYLAMMONIUM | -14.74 | -15.74 | 1.00 |
| HEXYLTRIMETHYLAMMONIUM | -13.85 | -13.88 | 0.03 |
| HYDROGEN ADIPATE | -12.4 | -12.85 | 0.45 |
| HYDROGEN FUMARATE | -12.4 | -11.77 | 0.63 |
| HYDROGEN GLUTARATE | -9.9 | -10.51 | 0.61 |
| HYDROGEN MALONATE | -8.6 | -8.34 | 0.26 |
| HYDROGEN OXALATE | -2.2 | -2.52 | 0.32 |
| HYDROGEN SUCCINATE | -7.1 | -7.71 | 0.61 |
| HYDROQUINONE | -6.85 | -8.40 | 1.55 |
| INDOLE | -44.3 | -35.78 | 8.52 |
| IODOBENZENE | -17.35 | -17.87 | 0.52 |
| L-HISTIDINE | -5.4 | -6.03 | 0.63 |
| L-MANDELIC ACID | -13 | -14.01 | 1.01 |
| L-METHIONINE | -5.4 | -6.61 | 1.21 |
| L-NORLEUCINE | -9.5 | -9.48 | 0.02 |
| L-NORVALINE | -6.2 | -6.94 | 0.74 |
| L-PHENYLALANINE | -6.5 | -7.81 | 1.31 |
| L-TRYPTOPHAN | -7.6 | -9.64 | 2.04 |
| L-TYROSINE | -8.2 | -9.25 | 1.05 |
| L-XYLOSE | -11.8 | -11.99 | 0.19 |
| MALEIC ACID | -8 | -14.19 | 6.19 |
| MALONIC ACID | -6.8 | -6.83 | 0.03 |
| M-CRESOL | -6.68 | -8.93 | 2.25 |
| METHANOL | 0.17 | -0.70 | 0.87 |
| M-METHYLCINNAMIC ACID | -23.58 | -22.62 | 0.96 |
| N-ACETYL-L-LEUCINAMIDE | -7.4 | -7.63 | 0.23 |
| NITROBENZENE | -17.81 | -16.36 | 1.45 |
| N-METHYLPHENETHYLAMMONIUM | -6.47 | -8.44 | 1.97 |
| OCTANEDIOATE | -12.6 | -12.70 | 0.10 |
| OCTANEDIOIC ACID | -18 | -17.83 | 0.17 |
| OCTANOATE | -16.9 | -17.32 | 0.42 |
| OCTANOIC ACID | -18.6 | -20.14 | 1.54 |
| OCTYLAMMONIUM | -19.27 | -19.75 | 0.48 |
| OXALIC ACID | -8 | -8.52 | 0.52 |
| P-CRESOL | -8.62 | -8.43 | 0.19 |
| PENTANOIC ACID | -15.56 | -15.16 | 0.40 |
| PENTYLAMINE | -15.2 | -14.36 | 0.84 |
| PENTYLAMMONIUM | -11.3 | -12.52 | 1.22 |
| PERCHLORATE | -7.5 | -7.29 | 0.21 |
| PERCHLORIC ACID | -9.2 | -9.06 | 0.14 |
| PHENETHYLAMMONIUM | -6.4 | -7.45 | 1.05 |
| PHENIRAMINE MALEATE | -10.4 | -11.44 | 1.04 |
| PHENOL | -6.17 | -10.30 | 4.13 |
| PHENYL ACETATE | -7.02 | -8.15 | 1.13 |
| PHENYL-BETA-D-GLUCOPYRANOSIDE | -5.6 | -6.56 | 0.96 |
| PIROXICAM | -9.6 | -10.61 | 1.01 |
| PROPIONIC ACID | -8.4 | -8.59 | 0.19 |
| PROPYLAMINE | -9.5 | -10.03 | 0.53 |
| P-XYLENE | -10.33 | -10.43 | 0.10 |
| PYRENE | -12.4 | -25.78 | 13.38 |
| PYRIDINE | -12.6 | -13.80 | 1.20 |
| QUINOLINE | -8.33 | -25.38 | 17.05 |
| RESORCINOL | -6.9 | -8.00 | 1.10 |
| SUCCINIC ACID | -11.4 | -11.85 | 0.45 |
| THIOCYANATE | -11.5 | -11.61 | 0.11 |
| TOLUENE | -12.27 | -13.97 | 1.70 |
| TRANS-1,2-CYCLOHEXANEDIOL | -7 | -7.35 | 0.35 |
| TRANS-2-HEXENOATE | -13.99 | -13.84 | 0.15 |
| TRANS-3-HEXENOATE | -12.6 | -13.61 | 1.01 |
| TRIETHYLAMINE | -9.9 | -10.87 | 0.97 |
| TRIMETHYLACETIC ACID | -6.9 | -6.99 | 0.09 |
| TYRAMINIUM | -5.55 | -7.65 | 2.10 |
| TYR-GLY-GLY-PHE-LEU | -7.2 | -7.64 | 0.44 |
| TYR-ILE-GLY-SER-ARG | -6.9 | -7.06 | 0.16 |

**Table 3.** Experimental and predicted free energy change for beta cyclodextrin.

| MOLECULE OF INTEREST | EXPERIMENTAL<br>(KJ/MOL) | PREDICTED<br>(KJ/MOL) | ABSOLUTE ERROR<br>(KJ/MOL) |
| --- | --- | --- | --- |
| 3-CHLOROPHENYL ACETATE | -13.93 | -12.80 | 1.13 |
| 3-ETHYLPHENOL | -14.84 | -15.78 | 0.94 |
| CYCLOOCTANOL | -18.84 | -18.76 | 0.08 |
| ETHYL BENZOATE` | -15.59 | -13.77 | 1.82 |
| KETOPROFEN | -16.31 | -16.34 | 0.03 |
| P-TOLYL ACETATE | -14.21 | -13.28 | 0.93 |
| THIANAPHTHENE | -18.44 | -17.30 | 1.14 |
| (1R,2R)-(-)-PSEUDOEPHEDRINE | -10.49 | -11.46 | 0.97 |
| (1R,2S)-(-)-EPHEDRINE | -10.84 | -9.73 | 1.11 |
| (1S,2R)-(+)-EPHEDRINE | -10.58 | -9.73 | 0.85 |
| (1S,2S)-(+)-PSEUDOEPHEDRINE | -11.33 | -11.46 | 0.13 |
| (2,5-DIMETHOXYPHENETHYL)AMMONIUM | -9.08 | -8.26 | 0.82 |
| (2-NAPHTHYLOXY)ACETIC ACID | -14.6 | -14.43 | 0.17 |
| (3,4-DIHYDROXYPHENETHYL)AMMONIUM | -8.58 | -9.03 | 0.45 |
| (3,4-DIMETHOXYPHENETHYL)AMMONIUM | -6.52 | -6.70 | 0.18 |
| (3-METHOXYPHENETHYL)AMMONIUM | -10.74 | -9.62 | 1.12 |
| (3-PHENYLPROPYL)AMMONIUM | -11.61 | -10.34 | 1.27 |
| (4-HYDROXYPHENETHYL)AMMONIUM | -10.53 | -10.23 | 0.30 |
| (4-METHOXYPHENETHYL)AMMONIUM | -11.23 | -10.45 | 0.78 |
| (4-METHYLPHENETHYL)AMMONIUM | -11.11 | -10.83 | 0.28 |
| (R)-(-)-2-BUTANOL | -6.4 | -6.23 | 0.17 |
| (R)-(-)-2-HEXANOL | -11.8 | -10.98 | 0.82 |
| (R)-(-)-2-PENTANOL | -8.7 | -8.43 | 0.27 |
| (R)-(-)-PHENYLEPHRINE | -19.44 | -16.19 | 3.25 |
| (S)-(+)-2-BUTANOL | -7.4 | -6.23 | 1.17 |
| (S)-(+)-2-PENTANOL | -8.6 | -8.43 | 0.17 |
| [2-(4-AMINOPHENYL)ETHYL]AMMONIUM | -8.54 | -7.60 | 0.94 |
| 1,2-ETHANEDIOL | 1.08 | 1.66 | 0.58 |
| 1,3-BUTANEDIOL | -23.3 | -18.03 | 5.27 |
| 1,3-PROPANEDIOL | -3.82 | -4.50 | 0.68 |
| 1,4-BUTANEDIOL | -3.65 | -3.65 | 0.00 |
| 1,4-DIBROMOBENZENE | -16.98 | -17.06 | 0.08 |
| 1,4-DIODOBENZENE | -18.12 | -18.07 | 0.05 |
| 1,5-PENTANEDIOL | -6.97 | -6.44 | 0.53 |
| 1,6-HEXANEDIOL | -9.65 | -9.29 | 0.36 |
| 1-ADAMANTANEACETATE | -24.69 | -23.56 | 1.13 |
| 1-ADAMANTANECARBOXYLATE | -25.7 | -26.22 | 0.52 |
| 1-ADAMANTANECARBOXYLIC ACID | -32.4 | -31.31 | 1.09 |
| 1-ADAMANTYLAMMONIUM | -28.7 | -28.01 | 0.69 |
| 1-AMINONAPHTHALENE | -14.1 | -13.37 | 0.73 |
| 1-BENZYLIMIDAZOLE | -14.92 | -13.91 | 1.01 |
| 1-BROMOMETHYLNAPHTHALENE | -16.6 | -16.47 | 0.13 |
| 1-BUTANOL | -6.99 | -6.12 | 0.87 |
| 1-BUTYLIMIDAZOLE | -12.5 | -12.31 | 0.19 |
| 1-CHLORO-4-NITROBENZENE | -12.27 | -11.96 | 0.31 |
| 1-CHLORONAPHTHALENE | -17.6 | -17.37 | 0.23 |
| 1-CYANONAPHTHALENE | -13.7 | -14.70 | 1.00 |
| 1-HEPTANOL | -16.27 | -16.11 | 0.16 |
| 1-HEXANOL | -13.31 | -11.94 | 1.37 |
| 1-METHOXYNAPHTHALENE | -17.9 | -16.41 | 1.49 |
| 1-METHYLCYCLOHEXANOL | -17.42 | -15.75 | 1.67 |
| 1-METHYLNAPHTHALENE | -16.2 | -17.31 | 1.11 |
| 1-NAPHTHALENEACETATE | -24.8 | -19.75 | 5.05 |
| 1-NAPHTHALENECARBOXAMIDE | -15 | -15.23 | 0.23 |
| 1-NAPHTHALENESULFONATE | -19.4 | -19.88 | 0.48 |
| 1-NAPHTHOL | -17.6 | -16.62 | 0.98 |
| 1-NAPHTHYL ACETATE | -12.5 | -13.78 | 1.28 |
| 1-NAPHTHYLALDEHYDE | -14.8 | -14.94 | 0.14 |
| 1-OCTANOL | -18.1 | -16.95 | 1.15 |
| 1-PENTANOL | -10.29 | -9.80 | 0.49 |
| 1-PROPANOL | -3.26 | -3.08 | 0.18 |
| 2-(DIMETHYLAMINO)-6-PROPIONYLNAPHTHALENE | -18.8 | -18.36 | 0.44 |
| 2,2-DIMETHYL-1-PROPANOL | -15.48 | -14.66 | 0.82 |
| 2,3,6-NAPHTHALENETRISULFONATE | -12.7 | -12.32 | 0.38 |
| 2,6-NAPHTHALENEDISULFONATE | -18.8 | -18.49 | 0.31 |
| 2,7-NAPHTHALENEDISULFONATE | -13.9 | -13.75 | 0.15 |
| 2-BUTANOL | -6.79 | -6.23 | 0.56 |
| 2-CHLOROPHENOL | -13.1 | -12.38 | 0.72 |
| 2-HEXANOL | -11.3 | -10.98 | 0.32 |
| 2-METHYL-1-BUTANOL | -11.87 | -10.73 | 1.14 |
| 2-METHYL-1-PROPANOL | -9.25 | -8.65 | 0.60 |
| 2-METHYL-2-BUTANOL | -10.9 | -10.39 | 0.51 |
| 2-METHYL-2-PENTANOL | -11.36 | -10.62 | 0.74 |
| 2-METHYL-2-PROPANOL | -9.59 | -9.02 | 0.57 |
| 2-METHYLCYCLOHEXANONE | -15.7 | -15.51 | 0.19 |
| 2-NAPHTHALENESULFONATE | -30.7 | -28.69 | 2.01 |
| 2-NAPHTHOATE | -14.6 | -17.96 | 3.36 |
| 2-NAPHTHOL | -15.4 | -15.26 | 0.14 |
| 2-NAPHTHYL ACETATE | -14.3 | -14.24 | 0.06 |
| 2-NORBORNANEACETATE | -20.5 | -20.59 | 0.09 |
| 2-OCTANOL | -17.87 | -16.26 | 1.61 |
| 2-PENTANOL | -8.51 | -8.43 | 0.08 |
| 2-PHENYLETHANOL | -12.27 | -11.69 | 0.58 |
| 2-PROPANOL | -3.6 | -2.68 | 0.92 |
| 3-(4-HYDROXYPHENYL)-1-PROPANOL | -17 | -16.89 | 0.11 |
| 3-(4-HYDROXYPHENYL)PROPIONATE | -14.11 | -13.72 | 0.39 |
| 3,3-DIMETHYL-2-BUTANOL | -15.7 | -15.50 | 0.20 |
| 3,5-DIBROMOPHENOL | -14.61 | -14.67 | 0.06 |
| 3,5-DICHLOROPHENOL | -11.82 | -11.91 | 0.09 |
| 3,5-DIMETHOXYPHENOL | -13.36 | -13.17 | 0.19 |
| 3-AMINOBENZOIC ACID | -10.3 | -11.51 | 1.21 |
| 3-BROMOPHENOL | -14.33 | -13.93 | 0.40 |
| 3-BROMOPHENYL ACETATE | -15.24 | -15.51 | 0.27 |
| 3-CHLOROPHENOL | -13.01 | -12.76 | 0.25 |
| 3-CYANOPHENYL ACETATE | -8.51 | -9.43 | 0.92 |
| 3-ETHOXYPHENOL | -13.41 | -14.37 | 0.96 |
| 3-ETHOXYPHENYL ACETATE | -14.21 | -13.73 | 0.48 |
| 3-ETHYLPHENYL ACETATE | -15.3 | -14.50 | 0.80 |
| 3-FLUOROPHENOL | -9.7 | -9.83 | 0.13 |
| 3-FLUOROPHENYL ACETATE | -10.9 | -10.37 | 0.53 |
| 3-HYDROXYACETOPHENONE | -11.76 | -12.57 | 0.81 |
| 3-HYDROXYBENZOIC ACID | -14 | -14.49 | 0.49 |
| 3'-HYDROXYPROPIOPHENONE | -14.9 | -13.60 | 1.30 |
| 3-IODOPHENOL | -16.73 | -16.71 | 0.02 |
| 3-IODOPHENYL ACETATE | -17.53 | -17.46 | 0.07 |
| 3-ISOBUTYLPHENOL | -24.03 | -22.65 | 1.38 |
| 3-ISOBUTYLPHENYL ACETATE | -21.86 | -20.28 | 1.58 |
| 3-ISOPROPYLPHENOL | -19.64 | -20.37 | 0.73 |
| 3-ISOPROPYLPHENYL ACETATE | -19.18 | -17.21 | 1.97 |
| 3-METHOXYPHENOL | -12.05 | -12.12 | 0.07 |
| 3-METHOXYPHENYLACETATE | -9.02 | -8.72 | 0.30 |
| 3-METHYL-1-BUTANOL | -12.84 | -12.01 | 0.83 |
| 3-METHYL-2-BUTANOL | -10.96 | -10.50 | 0.46 |
| 3-METHYL-3-PENTANOL | -12.27 | -11.58 | 0.69 |
| 3-METHYLBENZOIC ACID | -40.2 | -31.63 | 8.57 |
| 3-METHYLCYCLOHEXANOL | -16.66 | -16.55 | 0.11 |
| 3-METHYLPHENYL ACETATE | -12.61 | -11.63 | 0.98 |
| 3-METHYLPHENYLACETIC ACID | -6.1 | -9.16 | 3.06 |
| 3-N-BUTYLPHENOL | -21.46 | -20.48 | 0.98 |
| 3-N-BUTYLPHENYL ACETATE | -20.89 | -18.93 | 1.96 |
| 3-NITROANILINE | -9.1 | -9.26 | 0.16 |
| 3-NITROPHENOL | -14.9 | -14.25 | 0.65 |
| 3-N-PROPYLPHENYL ACETATE | -18.72 | -17.37 | 1.35 |
| 3-O-METHYLDOPAMINE | -5.9 | -7.22 | 1.32 |
| 3-PENTANOL | -7.71 | -7.01 | 0.70 |
| 3-PHENYLBUTANOATE | -14.76 | -15.16 | 0.40 |
| 3-PHENYLPROPIONATE | -12.27 | -12.52 | 0.25 |
| 3-PROPYLPHENOL | -18.72 | -17.87 | 0.85 |
| 3-SEC-BUTYLPHENOL | -23.18 | -21.53 | 1.65 |
| 3-TERT-BUTYLPHENOL | -25.18 | -24.96 | 0.22 |
| 4-AMINO BENZOIC ACID | -15.4 | -13.37 | 2.03 |
| 4-AMINO-N-METHYLPHthalimide | -11.8 | -11.38 | 0.42 |
| 4-BENZYLPIPERIDINE | -18.83 | -18.17 | 0.66 |
| 4-BROMOPHENOL | -15.12 | -15.07 | 0.05 |
| 4-BROMOPHENYL ACETATE | -15.3 | -14.85 | 0.45 |
| 4-CHLOROPHENOL | -14.92 | -14.63 | 0.29 |
| 4-CHLOROPHENYL ACETATE | -14.27 | -14.04 | 0.23 |
| 4-ETHOXYPHENOL | -13.3 | -13.79 | 0.49 |
| 4-ETHOXYPHENYL ACETATE | -14.5 | -14.36 | 0.14 |
| 4-ETHYLPHENOL | -15.37 | -15.77 | 0.40 |
| 4-ETHYLPHENYL ACETATE | -16.15 | -15.40 | 0.75 |
| 4-FLUOROPHENOL | -9.87 | -9.96 | 0.09 |
| 4-FLUOROPHENYL ACETATE | -12.05 | -11.73 | 0.32 |
| 4-HYDROXYACETOPHENONE | -12.44 | -11.93 | 0.51 |
| 4-HYDROXYBENZALDEHYDE | -9.99 | -10.34 | 0.35 |
| 4-HYDROXYBENZOIC ACID | -12.54 | -13.27 | 0.73 |
| 4-HYDROXYBENZYL ALCOHOL | -12.34 | -12.43 | 0.09 |
| 4-HYDROXYCINNAMIC ACID | -16.15 | -15.67 | 0.48 |
| 4-HYDROXYCOUMARIN | -13.1 | -12.96 | 0.14 |
| 4'-HYDROXYPROIOPHENONE | -15.01 | -14.37 | 0.64 |
| 4-IODOPHENOL | -17 | -16.83 | 0.17 |
| 4-IODOPHENYL ACETATE | -17.13 | -16.61 | 0.52 |
| 4-ISOPROPOXYPHENOL | -16.33 | -14.98 | 1.35 |
| 4-ISOPROPYLPHENOL | -20.43 | -18.64 | 1.79 |
| 4-ISOPROPYLPHENYL ACETATE | -16.44 | -17.02 | 0.58 |
| 4-METHOXYPHENOL | -12.61 | -13.06 | 0.45 |
| 4-METHOXYPHENYL ACETATE | -13.99 | -12.46 | 1.53 |
| 4-METHOXYPHENYLACETATE | -10.51 | -9.83 | 0.68 |
| 4-METHYL-2-PENTANOL | -11.64 | -10.84 | 0.80 |
| 4-N-AMYPHENOL | -23.92 | -23.40 | 0.52 |
| 4-N-AMYPHENYL ACETATE | -21.69 | -20.40 | 1.29 |
| 4-N-BUTYLPHENOL | -22.66 | -21.46 | 1.20 |
| 4-N-BUTYLPHENYL ACETATE | -20.66 | -19.51 | 1.15 |
| 4-NITROANILINE | -14.18 | -13.82 | 0.36 |
| 4-NITROBENZOIC ACID | -13.36 | -13.18 | 0.18 |
| 4-NITROPHENOL | -14.76 | -14.43 | 0.33 |
| 4-NITROPHENOLATE | -15 | -14.59 | 0.41 |
| 4-NITROPHENYL ACETATE | -12.16 | -11.72 | 0.44 |
| 4-NITROPHENYL-BETA-D-GALACTOSIDE | -10.1 | -8.99 | 1.11 |
| 4-NITROPHENYL-BETA-D-GLUCOSIDE | -8.8 | -8.99 | 0.19 |
| 4-N-PROPYLPHENYL ACETATE | -17.98 | -17.63 | 0.35 |
| 4-O-METHYLDOPAMINE | -10.71 | -9.41 | 1.30 |
| 4-PHENYLBUTANOATE | -15.06 | -14.52 | 0.54 |
| 4-PROPYLPHENOL | -20.26 | -19.57 | 0.69 |
| 4-SEC-BUTYLPHENOL | -23.86 | -23.37 | 0.49 |
| 4-TERT-AMYPHENOL | -26.83 | -26.57 | 0.26 |
| 4-TERT-BUTYLPHENOL | -26.03 | -25.05 | 0.98 |
| 4-TERT-BUTYLPHENYL ACETATE | -21.98 | -21.03 | 0.95 |
| 5-METHYLRESORCINOL | -9.8 | -10.10 | 0.30 |
| ACENOCOUMARIN | -14.7 | -14.51 | 0.19 |
| ACETALDEHYDE | 3.65 | 3.91 | 0.26 |
| ACETOANILIDE | -12.54 | -11.71 | 0.83 |
| ACETOHEXAMIDE | -16.77 | -16.14 | 0.63 |
| ACETONE | -2.4 | -1.96 | 0.44 |
| ACETONITRILE | 1.54 | 2.25 | 0.71 |
| ACETOPHENONE | -12.96 | -12.46 | 0.50 |
| ACRIDINE | -13.3 | -16.20 | 2.90 |
| ALLOBARBITAL | -11.32 | -10.92 | 0.40 |
| AMITRIPTYLIN | -25 | -23.99 | 1.01 |
| AMOBARBITAL | -17.53 | -16.35 | 1.18 |
| ANILINE | -9.13 | -8.46 | 0.67 |
| ANISOLE | -13.24 | -12.59 | 0.65 |
| ANTHRACENE | -18.9 | -19.37 | 0.47 |
| BARBITAL | -10.19 | -9.54 | 0.65 |
| BENDROFLUMETHIAZIDE | -10.83 | -9.89 | 0.94 |
| BENZ[A]ANTHRACENE | -20.2 | -19.40 | 0.80 |
| BENZALDEHYDE | -10.16 | -10.33 | 0.17 |
| BENZENE | -12.72 | -12.20 | 0.52 |
| BENZO[A]PYRENE | -19.1 | -21.39 | 2.29 |
| BENZOATE | -8.89 | -9.57 | 0.68 |
| BENZOIC ACID | -12.13 | -13.39 | 1.26 |
| BENZONITRILE | -12.74 | -11.69 | 1.05 |
| BENZOTHIAZOLE | -13.59 | -14.75 | 1.16 |
| BENZYL ALCOHOL | -9.76 | -9.68 | 0.08 |
| BENZYL CHLORIDE | -13.95 | -13.52 | 0.43 |
| BETAMETHASONE | -21.31 | -20.78 | 0.53 |
| BROMOBENZENE | -14.28 | -13.92 | 0.36 |
| BUTANOIC ACID | -8.62 | -9.43 | 0.81 |
| CARBAZOLE | -13.93 | -14.80 | 0.87 |
| CARBON TETRACHLORIDE | -12.56 | -11.88 | 0.68 |
| CARBUTAMIDE | -13.08 | -12.23 | 0.85 |
| CHENODEOXYCHOLIC ACID | -24.9 | -25.12 | 0.22 |
| CHLORAMINE-T | -18.1 | -17.47 | 0.63 |
| CHLORCYCLIZINE | -21 | -20.37 | 0.63 |
| CHLOROFORM | -8.16 | -7.53 | 0.63 |
| CHOLIC ACID | -19.97 | -19.88 | 0.09 |
| CINNARIZINE | -20.77 | -20.90 | 0.13 |
| CIS-1,2-CYCLOHEXANEDIOL | -13.9 | -11.53 | 2.37 |
| CIS-4-METHYLCYCLOHEXANOL | -18.07 | -17.21 | 0.86 |
| CORTICOSTERONE | -22 | -21.50 | 0.50 |
| CORTISONE | -19.1 | -18.92 | 0.18 |
| CORTISONE-21-ACETATE | -20.65 | -20.06 | 0.59 |
| CYCLOBARBITAL | -15.47 | -16.01 | 0.54 |
| CYCLOBUTANOL | -6.74 | -6.86 | 0.12 |
| CYCLOHEPTANOL | -19.08 | -17.76 | 1.32 |
| CYCLOHEXANECARBOXYLATE | -13.8 | -12.94 | 0.86 |
| CYCLOHEXANECARBOXYLIC ACID | -20.6 | -18.17 | 2.43 |
| CYCLOHEXANOL | -15.27 | -15.61 | 0.34 |
| CYCLOHEXANONE | -15.5 | -13.93 | 1.57 |
| CYCLOPENTANOL | -11.87 | -11.23 | 0.64 |
| DANSYLALANINE | -13 | -12.51 | 0.49 |
| DANSYL-L-ARGININE | -11.3 | -11.41 | 0.11 |
| DANSYL-L-ASPARAGINE | -11.7 | -11.42 | 0.28 |
| DANSYL-L-CYSTEIC ACID | -12.1 | -11.67 | 0.43 |
| DANSYL-L-GLUTAMINE | -12.1 | -11.50 | 0.60 |
| DANSYL-L-HISTIDINE | -12.6 | -12.54 | 0.06 |
| DANSYL-L-ISOLEUCINE | -13 | -12.74 | 0.26 |
| DANSYL-L-PHENYLALANINE | -10.9 | -11.85 | 0.95 |
| DANSYL-L-PROLINE | -13.4 | -13.37 | 0.03 |
| DANSYL-L-SERINE | -12.6 | -11.73 | 0.87 |
| DANSYL-L-TRYPTOPHAN | -14.6 | -12.73 | 1.87 |
| DANSYL-L-VALINE | -12.6 | -12.60 | 0.00 |
| DECANOATE | -21.8 | -20.05 | 1.75 |
| DECANOIC ACID | -22.9 | -21.70 | 1.20 |
| DEHYDROCHOLIC ACID | -19.29 | -19.41 | 0.12 |
| DEXAMETHASONE | -20.84 | -20.78 | 0.06 |
| D-GLUCOSE | -15 | -14.34 | 0.66 |
| DIBENZOFURAN | -16.95 | -16.74 | 0.21 |
| DIBENZOTHIOPHENE | -19.87 | -19.95 | 0.08 |
| DIDANSYL-L-LYSINE | -13 | -12.93 | 0.07 |
| DIETHYLAMINE | -7.77 | -6.98 | 0.79 |
| DIPHENYLPYRALINE | -20.4 | -20.18 | 0.22 |
| D-XYLOSE | -7 | -7.38 | 0.38 |
| ETHANOL | 0.17 | 0.53 | 0.36 |
| ETHYL BISCOUMACETATE | -20.4 | -20.36 | 0.04 |
| ETHYLBENZENE | -14.77 | -13.43 | 1.34 |
| FENBUFEN | -15.02 | -15.44 | 0.42 |
| FLUDIAZEPAM | -13.31 | -13.27 | 0.04 |
| FLUFENAMIC ACID | -17.7 | -16.70 | 1.00 |
| FLUOCINOLONE ACETONIDE | -19.85 | -19.37 | 0.48 |
| FLUOROBENZENE | -11.18 | -10.70 | 0.48 |
| FLURBIPROFEN | -21.07 | -20.02 | 1.05 |
| FUROSEMIDE | -10.19 | -10.08 | 0.11 |
| GRISEOFULVIN | -8.39 | -7.91 | 0.48 |
| HEPTANOATE | -14.2 | -14.00 | 0.20 |
| HEXANOATE | -10.44 | -8.93 | 1.51 |
| HEXANOIC ACID | -14.07 | -12.60 | 1.47 |
| HEXOBARBITAL | -17.6 | -16.72 | 0.88 |
| HEXYLAMMONIUM | -10.4 | -10.05 | 0.35 |
| HYDROCHLOROTHIAZIDE | -10.04 | -9.59 | 0.45 |
| HYDROCORTISONE | -20.58 | -20.65 | 0.07 |
| HYDROCORTISONE-17-BUTYRATE | -18.43 | -18.35 | 0.08 |
| HYDROQUINONE | -11.72 | -10.99 | 0.73 |
| IMIPRAMIN | -22.5 | -22.04 | 0.46 |
| IODOBENZENE | -16.71 | -16.54 | 0.17 |
| L-PHENYLALANINE | -7.2 | -7.05 | 0.15 |
| L-TRYPTOPHAN | -13.3 | -12.25 | 1.05 |
| L-TYROSINE | -8.7 | -8.67 | 0.03 |
| L-XYLOSE | -7.5 | -7.03 | 0.47 |
| MAPROTILIN | -21 | -21.94 | 0.94 |
| M-CRESOL | -11.3 | -11.14 | 0.16 |
| MECLOFENAMIC ACID | -15.27 | -15.20 | 0.07 |
| MEDAZEPAM | -13.72 | -14.14 | 0.42 |
| MEFENAMIC ACID | -14.24 | -13.60 | 0.64 |
| MENADION | -12.95 | -12.61 | 0.34 |
| MEPHOBARBITAL | -18.05 | -17.14 | 0.91 |
| METHANOL | 2.8 | 3.09 | 0.29 |
| METHYL BENZOATE | -14.28 | -14.38 | 0.10 |
| M-METHYLCINNAMIC ACID | -16.75 | -17.53 | 0.78 |
| N,N-DIMETHYLANILINE | -13.48 | -12.71 | 0.77 |
| N-ACETYL-1-AMINONAPHTHALENE | -12.1 | -12.60 | 0.50 |
| NAPHTHALENE | -16.2 | -16.49 | 0.29 |
| N-ETHYLANILINE | -13.33 | -12.77 | 0.56 |
| NIFLUMIC ACID | -15.5 | -15.39 | 0.11 |
| NIMETAZEPAM | -9.89 | -9.89 | 0.00 |
| NITRAZEPAM | -11.26 | -10.77 | 0.49 |
| NITROBENZENE | -11.64 | -11.08 | 0.56 |
| N-METHYLANILINE | -12.07 | -12.03 | 0.04 |
| N-METHYLPHENETHYLAMMONIUM | -8.3 | -8.63 | 0.33 |
| N-PHENYLANTHRANILIC ACID | -16.53 | -15.67 | 0.86 |
| OCTANOATE | -17.7 | -16.73 | 0.97 |
| OCTANOIC ACID | -15.7 | -17.59 | 1.89 |
| OCTYLAMMONIUM | -15 | -14.06 | 0.94 |
| PARAMETHASONE | -19.44 | -19.18 | 0.26 |
| P-CRESOL | -13.68 | -14.32 | 0.64 |
| PENTANOIC ACID | -11.16 | -11.26 | 0.10 |
| PENTOBARBITAL | -17.22 | -15.45 | 1.77 |
| PHENANTHRIDINE | -14.67 | -16.18 | 1.51 |
| PHENAZINE | -13.76 | -18.77 | 5.01 |
| PHENETHYLAMMONIUM | -7.9 | -7.68 | 0.22 |
| PHENETOLE | -14.2 | -13.06 | 1.14 |
| PHENIRAMINE MALEATE | -14.2 | -14.22 | 0.02 |
| PHENOBARBITAL | -18.37 | -16.73 | 1.64 |
| PHENOL | -11.28 | -11.20 | 0.08 |
| PHENOTHIAZINE | -15.59 | -15.83 | 0.24 |
| PHENOXAZINE | -15.36 | -15.34 | 0.02 |
| PHENPROCOUMON | -16.2 | -16.56 | 0.36 |
| PHENYL ACETATE | -11.99 | -11.80 | 0.19 |
| PHENYLACETYLENE | -13.48 | -13.77 | 0.29 |
| PIROXICAM | -11.2 | -10.81 | 0.39 |
| PREDNISOLONE | -20.3 | -20.07 | 0.23 |
| PROGABIDE | -14.48 | -14.18 | 0.30 |
| PROPANOLOL | -12.7 | -12.68 | 0.02 |
| PROSTAGLANDIN E2 | -14.1 | -13.79 | 0.31 |
| P-XYLENE | -13.58 | -12.75 | 0.83 |
| PYRENE | -9.1 | -20.80 | 11.70 |
| PYRILAMINE MALEATE | -16.3 | -15.04 | 1.26 |
| QUINOLINE | -12.1 | -15.63 | 3.53 |
| SPIRONOLACTONE | -25.34 | -25.22 | 0.12 |
| SULFADIAZINE | -15.7 | -14.45 | 1.25 |
| SULFADIMETHOXINE | -12.89 | -12.80 | 0.09 |
| SULFAETHIDOLE | -18 | -17.44 | 0.56 |
| SULFAGUANIDINE | -14.6 | -14.41 | 0.19 |
| SULFAMERAZINE | -11.8 | -12.59 | 0.79 |
| SULFAMETHAZINE | -11.8 | -10.56 | 1.24 |
| SULFAMETHIZOLE | -16.8 | -15.81 | 0.99 |
| SULFAMETHOMIDINE | -13.31 | -12.54 | 0.77 |
| SULFAMONOMETHOXINE | -14.16 | -12.97 | 1.19 |
| SULFANILAMIDE | -15.79 | -15.23 | 0.56 |
| SULFAPHENAZOLE | -13.42 | -13.19 | 0.23 |
| SULFAPYRIDINE | -15.43 | -15.98 | 0.55 |
| SULFATHIAZOLE | -19.24 | -18.47 | 0.77 |
| SULFISOMIDINE | -12.02 | -11.88 | 0.14 |
| SULFISOXAZOLE | -13.25 | -13.75 | 0.50 |
| TETRAHYDROFURAN | -8.39 | -7.97 | 0.42 |
| THENYLDIAMINE | -13.2 | -12.57 | 0.63 |
| THIANTHRENE | -20.38 | -20.35 | 0.03 |
| THIOPENTAL | -18.71 | -17.91 | 0.80 |
| TOLNAFTATE | -21.9 | -21.62 | 0.28 |
| TOLUENE | -11.93 | -11.62 | 0.31 |
| TRANS-1,2-CYCLOHEXANEDIOL | -11.4 | -11.53 | 0.13 |
| TRIAMCINOLONE | -19.27 | -19.11 | 0.16 |
| TRIAMCINOLONE ACETONIDE | -20.03 | -19.69 | 0.34 |
| TROPICAMIDE | -15 | -13.92 | 1.08 |
| TYR-GLY-GLY-PHE-LEU | -11.4 | -10.88 | 0.52 |
| TYR-ILE-GLY-SER-ARG | -13.2 | -12.61 | 0.59 |
| URSODEOXYCHOLIC ACID | -25.71 | -25.12 | 0.59 |
| VALETHAMATE BROMIDE | -24 | -23.36 | 0.64 |
| WARFARIN | -16.3 | -15.66 | 0.64 |
| XANTHENE | -15.47 | -15.54 | 0.07 |

**Table 4.** Experimental and predicted free energy change for gamma cyclodextrin.

| MOLECULE OF INTEREST | EXPERIMENTAL<br>(KJ/MOL) | PREDICTED<br>(KJ/MOL) | ABSOLUTE ERROR<br>(KJ/MOL) |
| --- | --- | --- | --- |
| 4-AMINO-1-NAPHTHALENESULFONATE | -7.5 | -6.53 | 0.97 |
| ANTHRACENE | -13.4 | -10.90 | 2.50 |
| BENZ[A]ANTHRACENE | -15.9 | -15.45 | 0.45 |
| BENZENE | -5.47 | -7.76 | 2.29 |
| 2-CHLORO-4-[(4-HYDROXYPHENYL)AZO]BENZOATE | -24.55 | -23.63 | 0.92 |
| 4-CHLORO-3-[(4-HYDROXYPHENYL)AZO]BENZOATE | -23.75 | -22.78 | 0.97 |
| FLURBIPROFEN | -19.9 | -18.98 | 0.92 |
| 3-[(4-HYDROXYPHENYL)AZO]BENZOATE | -22.49 | -22.03 | 0.46 |
| 4-[(4-HYDROXYPHENYL)AZO]BENZOATE | -23.57 | -22.11 | 1.46 |
| 2,7-NAPHTHALENEDISULFONATE | -14.7 | -13.77 | 0.93 |
| 2-NAPHTHALENESULFONATE | -6.6 | -6.52 | 0.08 |
| NIFLUMIC ACID | -13.9 | -12.99 | 0.91 |
| PIROXICAM | -9.8 | -8.68 | 1.12 |
| PYRENE | -17.4 | -11.44 | 5.96 |
| RESORCINOL | -7.7 | -6.93 | 0.77 |

**Figure 4.**
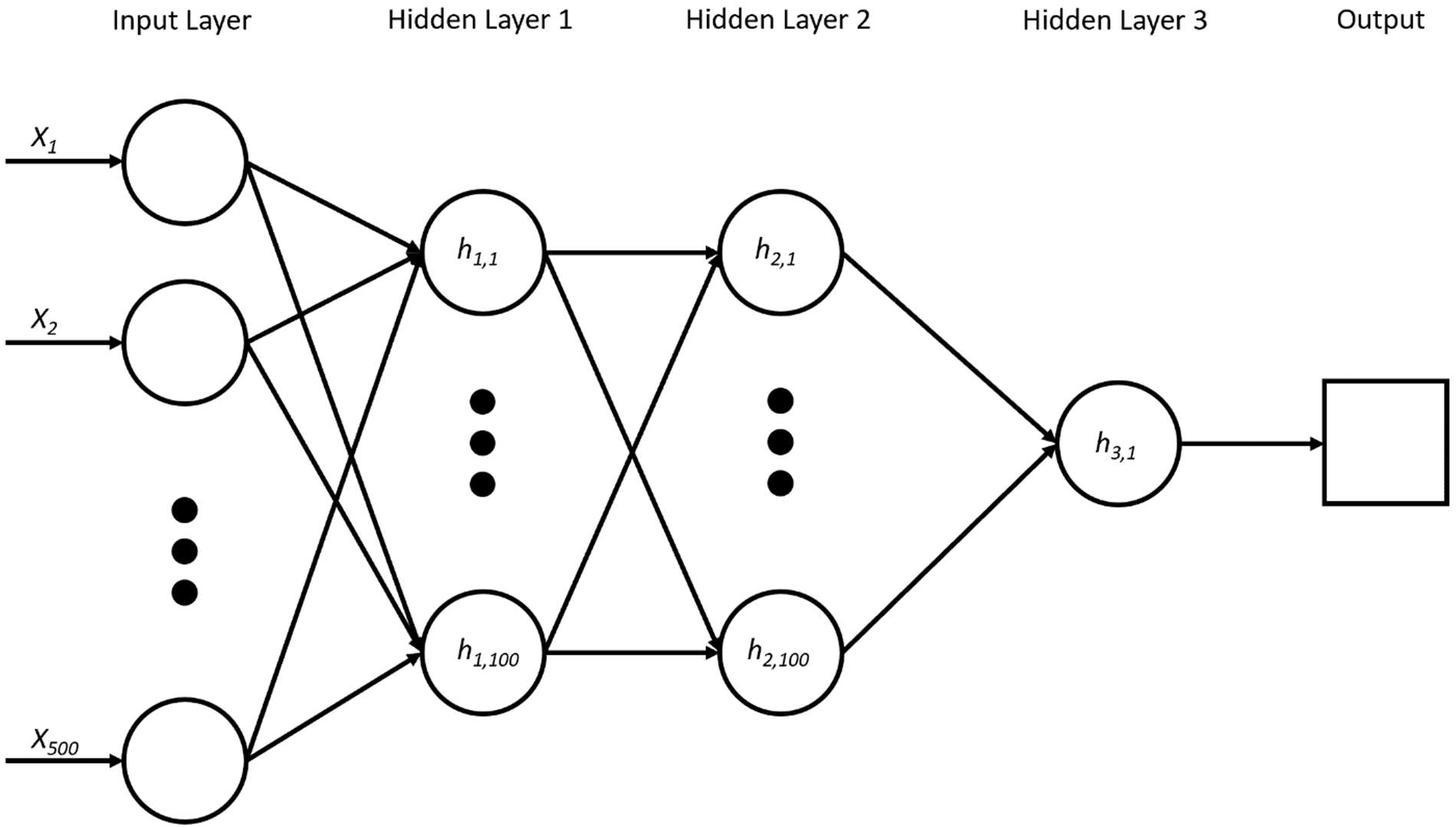
Optimal network design determined via model training.

**Figure 5.**
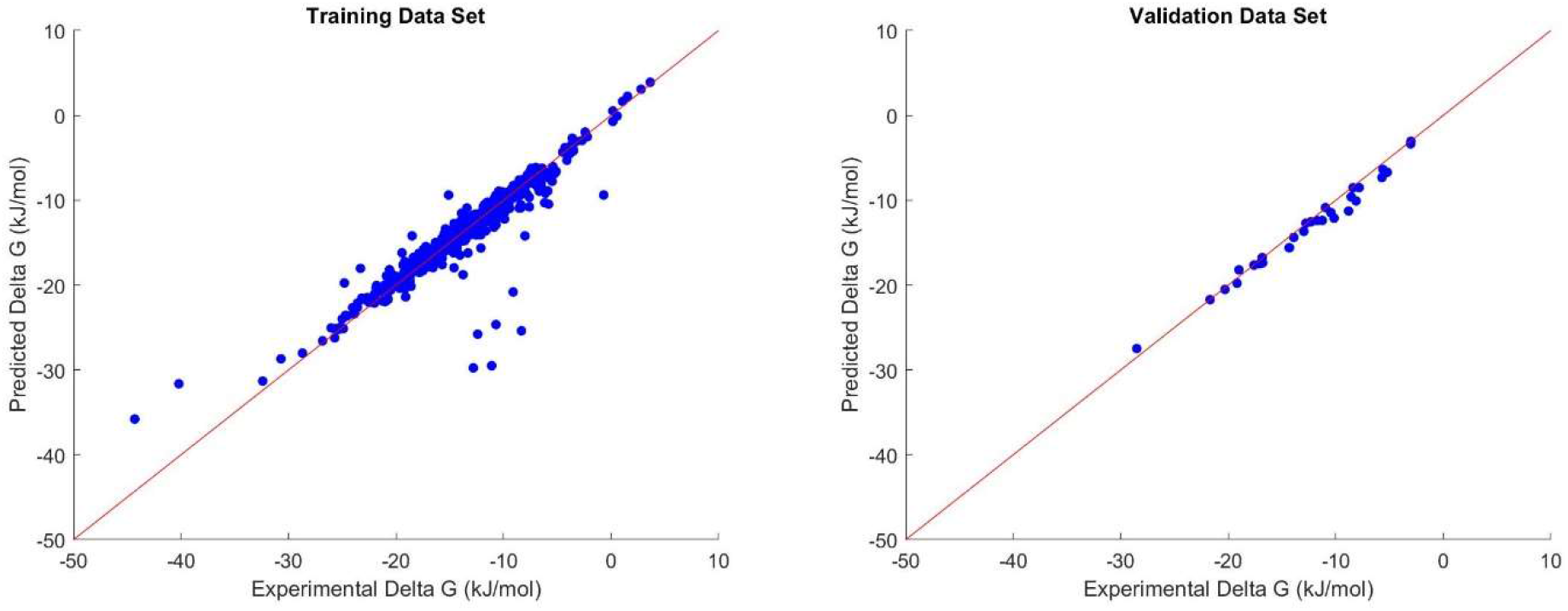
Graphical representation of the correlation between experimental and predicted values in data set used to train the model (A, left) and to validate the model (B, right). High linearity is representative of high correlation between predicted and experimental binding affinities.

Molecules that had largest error in predicted values, seen in Figure 5A, were aromatic compounds comprised of one or more benzene rings (i.e. pyrene, anthracene). It is interesting to note that in all three polymer cases, benzene and most benzene derivatives had low error values. This error may be caused by the removal of an important feature in the initial feature cleaning steps or due to the small number of similar molecules. Using a different QSAR descriptor calculator (i.e. Dragon) capable of calculating a greater number of descriptors may be minimize this error. An increase in samples used in the training set of large aromatic compounds may give the model enough examples to minimize this error as well.

## 4. Conclusion

The proposed model accurately predicted the binding energy between three different model polymers made from different cyclodextrins, and a large variety of small molecule drugs under physiological relevant conditions in both the training and test sets. While this first model is most useful in predicting potential interactions between CD polymers and a broad variety of small molecule drugs in biologically relevant environments there are as yet limited biomedical applications of cyclodextrins. Nevertheless, this model will help predict and optimize drug/polymer interactions in a high throughput manner. As one of the first models of its type, the initial use might be limited. But future versions of this model we would expect to be applied to much broader categories of polymer, provided there is enough impetus to generate the physical data needed to generate these models. We hope this work with inspire both future biomaterials models as well as wet-research, with the final to more rapidly aide in screening, and development of biomedical implants.

